# “A transient heritable memory regulates HIV reactivation from latency”

**DOI:** 10.1101/2020.07.02.185215

**Authors:** Yiyang Lu, Abhyudai Singh, Roy D. Dar

## Abstract

Reactivation of human immunodeficiency virus 1 (HIV-1) from latently infected T cells is a critical barrier to successfully eradicate HIV-1 from patients. Latency models in Jurkat T-cells reveal that individual cells reactivate in an all-or-none fashion when exposed to latency reversal agents (LRAs). Remarkably, this heterogeneity arises within a clonal cell population of genetically identical cells containing a single copy of the provirus integrated at the same genomic site. To characterize these single-cell responses, we leverage the classical Luria-Delbrück fluctuation test where single cells are isolated from a clonal population, and exposed to LRAs after a period of colony expansion. If cellular responses are purely random, then the fraction of reactivating cells should have minimal colony-to-colony fluctuations given the large number of cells present after weeks of colony growth. In contrast, data shows considerable colony-to-colony fluctuations with the fraction of reactivating cells following a skewed distribution. Systematic measurements of fluctuations over time in combination with mathematical modeling uncovers the existence of a heritable memory that regulates HIV-1 reactivation, where single cells are in a LRA-responsive state for a few weeks before switching back to an irresponsive state. These results have enormous implications for designing therapies to purge the latent reservoir and illustrate the utility of fluctuation-based assays to uncover hidden transient cellular states underlying phenotypic heterogeneity.

## Introduction

With over 37 million infected individuals, human immunodeficiency virus 1 (HIV-1) remains a global epidemic of unprecedented proportions. Upon infection with HIV-1, CD4+ T-cells may progress into a quiescent state called latency where they do not express viral genes and are capable of evading drug treatment. Latently infected cells can reactivate later and reinitiate active replication, causing a resurgence in viral levels if the patient goes off antiretroviral treatment (Fig. 1A). Actively replicating virus hijacks the host-cell resources and machinery to create hundreds of viral progeny, lyse the cell, and spread to uninfected bystander cells (1, 2). Thus HIV-1 latency remains the major barrier to a cure (3-7). Several strategies exist to control and clear the latent reservoir of HIV-1 infected cells (3-5). Much attention and research has focused on targeting the latent pool of cells with a small molecule treatment called the “shock and kill” strategy (8-10). In this treatment, cells are reactivated from latency into an active state where they can then undergo killing by cytotoxic T-cells and subsequently were cleared from the patient (Fig. 1A) (3). Unfortunately, “shock and kill” falls short due to incomplete reactivation and killing of the latent reservoir (11, 12). Further, latency reversal agents (LRAs) have been shown to impair cytotoxic T-cell function (13-15), and clinical trials with histone deacetylase inhibitors (HDACis) and disulfiram, two classes of candidate LRAs, produced no major reduction in the size of the latent HIV-1 reservoir in patients on antiretroviral therapy (ART) (10). Alternatively, latency promoting agents (LPAs) have also been discovered that silence transcription of latent HIV-1 (16-18), which points to the alternative “block and lock” strategy aiming to stabilize and prolong HIV-1 latency indefinitely.

**Figure 1.**
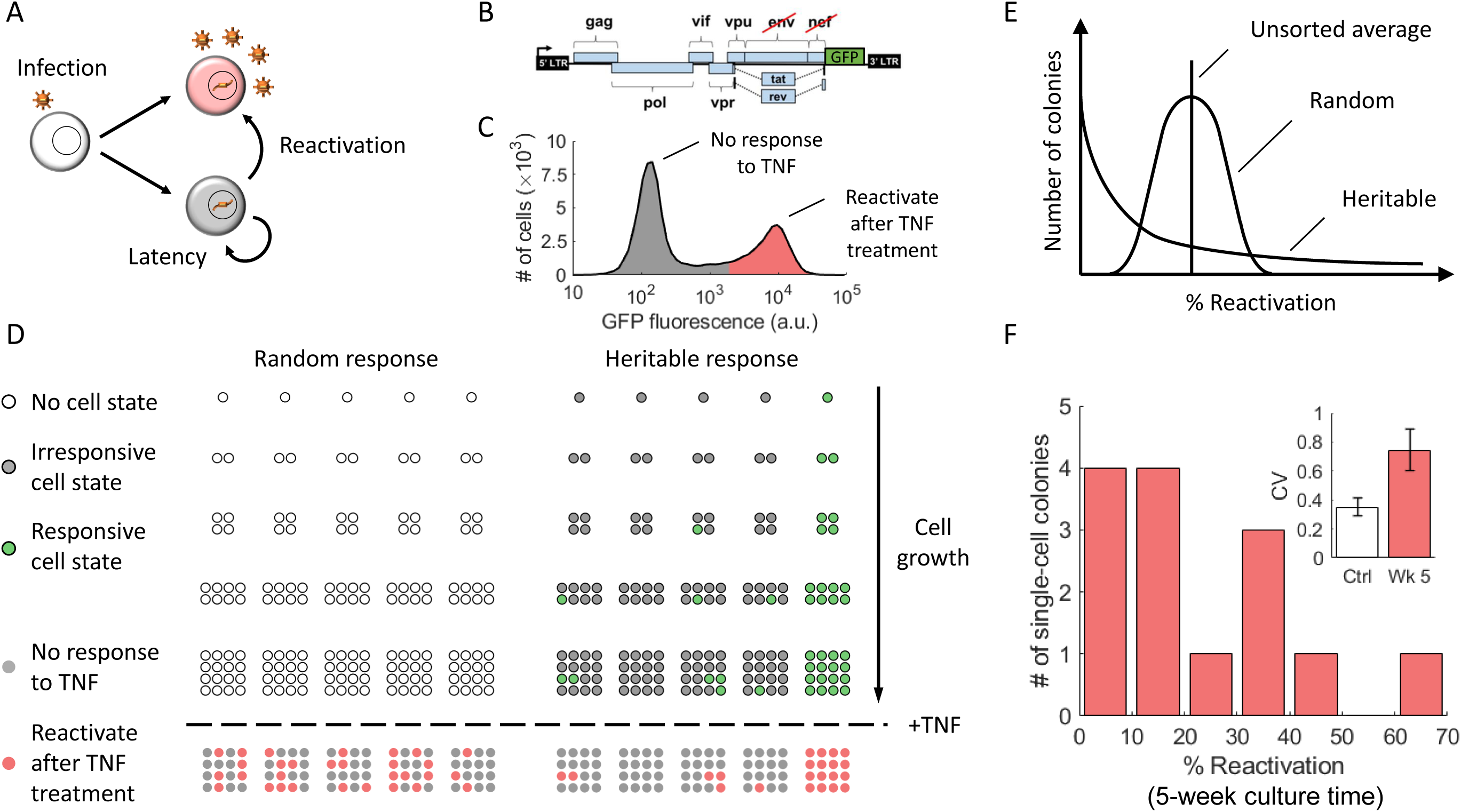
Heritable versus random reactivation response of HIV-1 latency to an external stimulus. **(A)** HIV-1 infection results in two phenotypes: active replication (red) where the infected cells produce new virions, and latent infection (gray) where infected cells do not produce virions. Latently infected cells can reactivate and initiate virion production through cellular and environmental fluctuations. **(B)** The cell line used in our investigation, the Jurkat latency model (JLat), is latently infected with a full-length HIV-1 gene circuit, with a deletion of *env* reading frame and a replacement of *nef* reading frame with a GFP element. **(C)** Histogram of single-cell GFP fluorescence (measured with flow cytometry) from a Jlat 9.2 bulk culture, after a 24-hour tumor necrosis factor α (TNF-α) treatment. TNF-α is a potent activator of the HIV-1 LTR promoter and was administered at 10 ng/ml concentration. The histogram shows that there is a distinct bimodal distribution of response to TNF-α perturbation within a clonal population of JLat. **(D)** We propose the following Luria-Delbrück experimental design to investigate if single-cell JLat response to TNF-α perturbation are random or heritable. A clonal population of JLat 9.2 were sorted into single cells using FACS and cultured individually into colonies. The colonies were subject to 10 ng/ml of TNF-α and the percentage of reactivated cells were measured using flow cytometry after 24 hours of treatment. If the JLat responds to TNF-α randomly, their responsiveness to TNF-α is determined at the time of TNF-α addition and thus would see low colony-to-colony variation. In contrast, if the response is heritable, then colony behaviors would be influenced by the parent cell resulting in large variations between colonies. **(E)** The distribution of reactivation percentage would be centered around the unsorted average if the response is random, while the heritable model will result in a wide, skewed distribution for reactivation percentage across colonies. **(F)** Histogram of reactivation percentage of 14 different JLat 9.2 colonies after a 24-hour TNF-α treatment. Colonies were cultured for 5 weeks after FACS and were grown from single cells sorted from a bulk JLat 9.2 clonal population. The resulting distribution coincides with the heritable response prediction, and confirms that JLat cells possess heritable memory of responsiveness to TNF-α perturbation. Inset: The noise in percent reactivation (quantified using the coefficient of variation CV) is significantly higher than the control noise floor as measured by fluctuations in percent reactivation across unsorted JLat populations.

Full control of the latent cell reservoir with small molecules, either by reactivation or silencing, requires a precise understanding of whether latent cells undergoing clonal expansion (19) have memory of their parent cell’s responsiveness to treatments. In this context one can envision two different mechanisms of HIV-1 reactivation: 1) A random model where individual cells reactivate purely stochastically owing to noise in post-exposure signaling and viral gene expression (20-24); 2) An alternative heritable model where reactivation is a deterministic function of an underlying cell state (for example, HIV-1 promoter’s epigenetic signature) just prior to LRA exposure. Despite its importance for understanding reactivation dynamics of the latent cell population, it is currently unknown which of the two cases, random response versus non-random heritable response, occurs in T-cells latently infected with HIV-1.

To discriminate between the random vs. heritable model we apply the Luria-Delbrück fluctuation test on a cell line model of HIV-1 latency (Fig. 1B-1C). More specifically, single cells are isolated from a clonal cell population having a single copy of the latent provirus at a unique integration site. Each single cell is expanded into a colony, and colony-to-colony fluctuations in the fraction of reactivating cells are quantified after LRA exposure (Fig. 1D). In the random model, each individual cell has the same probability 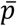 of reactivation, in which case the number of reactivating cells follows a Binomial distribution. A straightforward calculation shows that for the random model, the colony-to-colony fluctuations in the fraction of reactivating cells (as quantified using the coefficient of variation) is 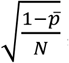, where *N* is the number of cells in the colony. Thus, for a sufficiently large *N*, the variations across colonies will be minimal, with each colony reactivating to the same frequency as the original clonal population. In the heritable model, the state of the single cell is propagated across generations, i.e., a more LRA responsive single cell leads to a more responsive colony. This memory of the starting cell state in the expanded population drives considerable colony-to-colony fluctuations in the heritable model (Fig. 1E). We develop a mathematical modeling framework to predict colony-to-colony differences for a transiently heritable cell state, and systematically couple them to experimentally measured fluctuations to elucidate mechanisms regulating HIV-1 reactivation.

## Results

To test which of the two types of response memory occurs in a latent cell population we utilized and cultured a known Jurkat latency model of HIV-1 (JLat, Fig. 1B) (25). This model consists of a Jurkat T-cell population latently infected with a full-length HIV-1 construct. The construct contains a deletion of *env* and a replacement of *nef* reading frame with GFP. JLat 9.2, a clonal cell population with a single HIV-1 integration site, was selected for our experiments (25). Tumor necrosis factor alpha (TNF-α) potently activates transcription of the HIV-1 LTR promoter through its NF-κB binding sites (26), and has been used for latency reversal assays on JLat cell lines in previous literature at 10 ng/ml (17, 27, 28). A typical response to TNF-α addition after 24h treatment is shown in Figure 1C. Here the bimodal distribution of identical cells shows a lower GFP peak of a cell subpopulation irresponsive to TNF-α and a higher GFP peak of cells that reactivate in response to TNF-α treatment (Fig. 1C) (1, 2). The percentage of reactivated cells measured using flow cytometry is quantified by the number of cells that turn on past a gating threshold divided by the total number of cells (Fig. S2), and is ≈ 20% for JLat 9.2 at 10 ng/ml TNF-α induction.

To perform the Luria-Delbrück fluctuation test, we used fluorescence activated cell sorting (FACS) to sort single cells into 96-well plates and grow out JLat single-cell colonies (Methods, Fig. S1). Of the cells that survived and expanded, 14 JLat 9.2 single-cell colonies were grown out over 5 weeks and subsequently treated with TNF-α for 24 hours to determine the fraction of reactivating cells (% reactivation). Intriguingly, the data reveals a skewed distribution for the % reactivation across colonies (Fig. 1F) with a high coefficient of variation (CV) of ≈ 0.75. While the parent JLat 9.2 population shows a 20% reactivation, 4 out of 14 single-cell colonies reactivate less than 10%, and 2 colonies reactivate greater than 40%. The skewed distribution observed at week 5 closely resembles one expected from a non-random heritable response model (Fig. 1E-1F). Next, we quantified the technical background noise in this assay, by measuring the % reactivation in unsorted JLat 9.2 populations over an eight-week period. The CV of week-to-week fluctuations in these unsorted populations was determined to be ≈ 0.35, and represents the control noise floor of the fluctuation test. The significantly higher fluctuations in the % reactivation across single-cell colonies (CV ≈ 0.75) as compared to the control noise floor (Fig. 1F; inset, p < 0.005) rules out the random response model, but is consistent with a heritable cell state regulating cellular responsiveness to TNF-α.

The cell outgrowth assay followed by activator treatment reveals that JLat 9.2 cells exhibit non-random heritable memory with respect to TNF-α activation. If the cell state dictating HIV-1 reactivation is transient, then the fluctuations in the % reactivation should decay over time and reach the noise floor. To determine the timescale of this noise relaxation, we measured TNF-α response across 16 single-cell colonies (14 colonies at week 5 plus 2 additional colonies that grew at week 7) every 2 weeks starting from week 5 (Fig. 2A). The reactivation level of the parental unsorted JLat 9.2 cell population was acquired to compare colony reactivation with unsorted reactivation. In brief, the unsorted JLat 9.2 population was cultured in duplicate and cells were measured by flow cytometry every two weeks after 24-hour TNF-α treatments. For each time point, the mean of 2 JLat 9.2 unsorted populations was acquired. Unsorted % reactivation drifted from about 32% in week 5 to about 12% in week 13 (black dots, Fig. 2A). The difference between each colony-wise % reactivation and the unsorted (Δ% Reactivation) decreased as culture time went on (Fig. 2B and 2C). As expected for a transiently heritable cell state, the colony-to-colony fluctuations in the % reactivation attenuate over time, starting from a wide, skewed distribution at week 5, to a narrow distribution at week 13 (Fig. 2C). The CV of the % reactivation at week 5 and 7 is significantly higher than the control noise floor, but drops to the noise floor starting from week 9. As also observed in the unsorted populations, the mean % reactivation from week 11 and week 13 are significantly lower than the control mean (p < 0.05 and p < 0.01 respectively), while weeks 5 to 9 do not show significant changes compared to the control (Fig. 2D).

**Figure 2.**
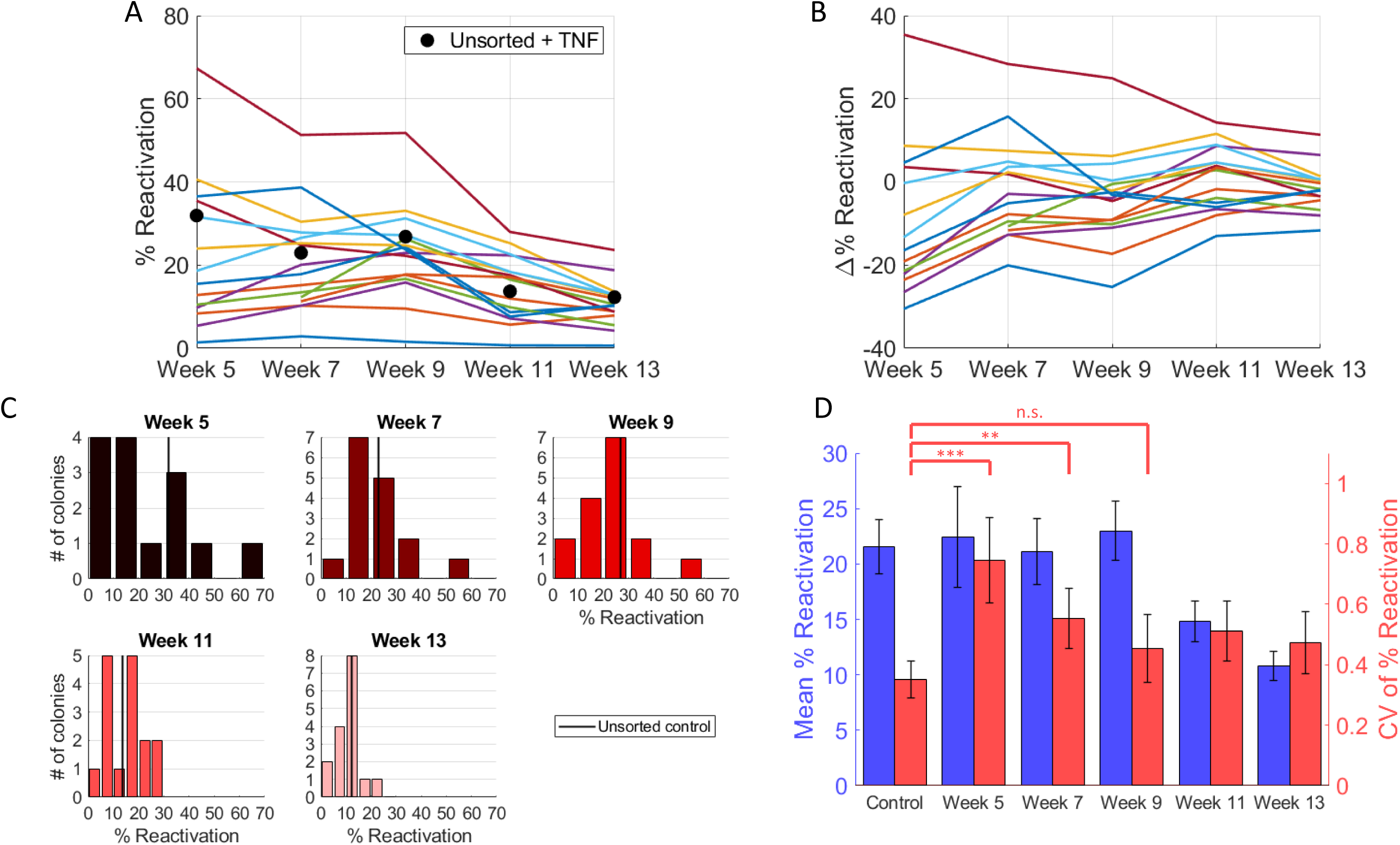
Experimental results from long-term culturing of single-cell JLat colonies reveal a transient heritable response to TNF-α perturbation. **(A)** Reactivation percentage of each colony across weeks 5 to 13 after TNF-α exposure of 24 hours, tested biweekly. Two colonies did not reach experimentally feasible concentrations at week 5, and were included starting from week 7. Black dots represent the control values, measured by exposing the unsorted culture to the same TNF-α treatment for 24 hours. The colonies gradually moved closer towards the unsorted control as culturing time progressed. This trend is more clearly represented in panel **(B)** where the difference in reactivation percentage between each colony and the control, Δ% Reactivation, is visualized. The approach towards unsorted control is demonstrated by the bundling of colony-wise response around 0 at later time points. **(C)** Histogram of reactivation percentage of single-cell colonies from each week’s measurement. Week 5 has a total of 14 colonies, while the rest weeks have 16. Unsorted control average is shown as black vertical lines. Bin size for weeks 5, 7 and 9 was set to 10, while for weeks 11 and 13 it was set to 5 for better resolution. The histograms also show the convergence of colony-wise behavior towards the unsorted control. **(D)** Bar plot of the colony-wise mean reactivation percentage (blue) and colony-wise mean CV (coefficient of variation squared) of reactivation percentage (red). Mean % reactivation values showed an overall decreasing trend as culture time went on, while mean CV of % reactivation decreased for the first 5 weeks before reaching a plateau in weeks 11 and 13. The control value is a pooled average of all unsorted +TNF-α treatment values across the 5 time points. Bar heights and error bars represent bootstrapped mean and standard error. Levels of significance are indicated as n.s. (p ≥ 0.1), one star (*, 0.1 > p ≥ 0.05), two stars (**, 0.05 > p ≥ 0.01), or three stars (***, p < 0.01). p-values were calculated using bootstrapping. See Methods for details on statistics.

To systematically characterize the hidden memory regulating HIV reactivation from the fluctuation test data, we develop a mathematical model of clonal expansion where single cells switch between an irresponsive and responsive state (Fig. 3A). The states are transiently heritable, i.e., the state is inherited for some generations before it is lost. Let *p*(*t*) denote the reactivation probability of an individual cell at time *t*, and is assumed to switch between two values: *p*(*t*) = 0 (irresponsive state) and *p*(*t*) = 1 (fully responsive state). Cells in the irresponsive state become responsive with rate *k*, and responsive cells become irresponsive with rate γ. In the stochastic formulation of the model, the time spent in responsive and irresponsive states is exponentially distributed with means 1/γ and *1/k*, respectively. These switching rates yield the following steady-state mean and noise levels of *p*(*t*) (see Material and Methods for details on model description and analysis)

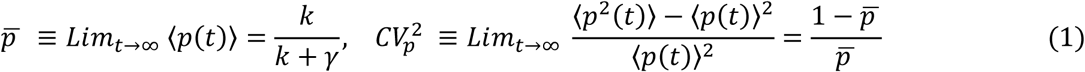

where the angular brackets ⟨*p*⟩ denote the expected value, and 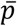 is the fraction of responsive cells in the original unsorted population.

**Figure 3.**
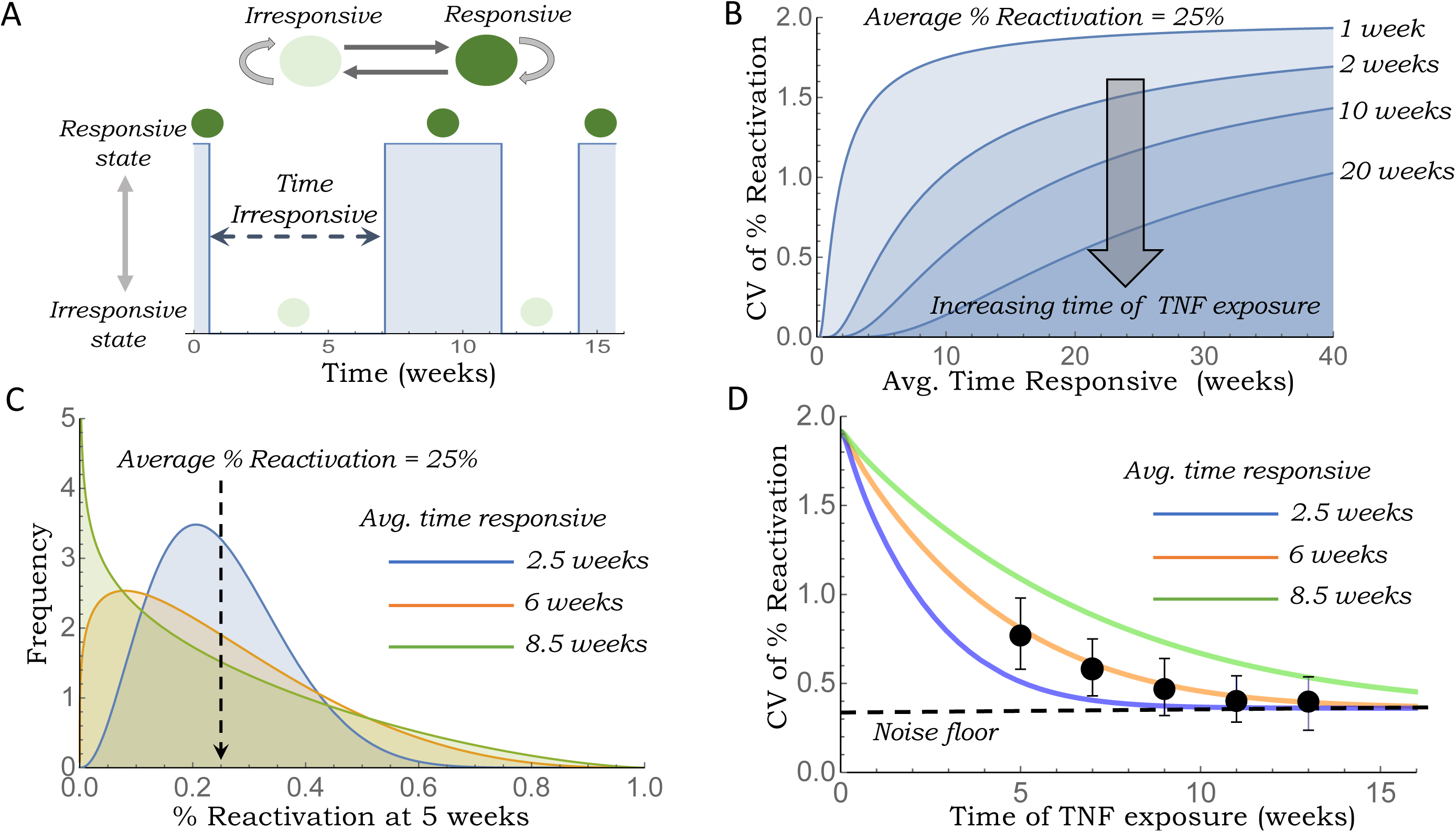
A transient cell state governs HIV-1 exit from latency. **(A)** Schematic of the stochastic model where prior to TNF-α exposure, individual cells reversibly switch between a TNF-α irresponsive and a responsive state. Self arrows represent proliferation of cells with the mother cell state being inherited by daughters. A sample realization of cells switching states over time is shown below. **(B)** The predicted colony-to-colony fluctuations in the fraction of reactivating cells (as measured by the coefficient of variation CV) increases for longer periods in the responsive state assuming a fixed *25%* reactivation averaged across colonies. Plots are shown for different times of TNF-α exposure. **(C)** Predicted distribution of the fraction of reactivating cells across colonies after 5 weeks of growth for different average times in the responsive state. Distributions are obtained by using a beta distribution with an average 25% reactivation and noise as given by equation (2). **(D)** Measured colony-to-colony fluctuations (with error bars showing standard error) starting from 5 to 13 weeks (black circles corresponding to data from Figure 2D). The fluctuations are considerably high at the initial 5-week timepoint (CV *=* 75%) and then gradually decrease to the noise floor. Note that the 11 and 13-week timepoints have been corrected for the mean as discussed in the main text. Model predicted CV as per equation (4) are shown assuming a noise floor *CV*_*NF*_ *= 0*.*35* for different average times in the responsive state.

Having defined stochastic transitions between the two cell states, we next model the clonal expansion in the Luria-Delbrück experiment. A single cell is chosen from the original population at time *t = 0*, and this initial cell is either in the responsive state with probability 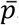, or in the irresponsive state with probability 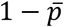. Starting from a single cell, colony size increases exponentially over time with cell-doubling times assumed to be the same irrespective of the underlying state. As the lineage expands, single cells reversibly switch between states, and the mother cell state is inherited by both daughters upon mitosis. Considering exposure to TNF-α at time *t*, our analysis predicts the CV of the colony-to-colony fluctuations in % reactivation

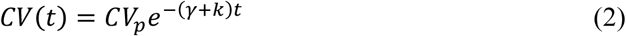

to decay exponentially with rate *k* + γ (see Material and Methods for details). Note that if reactivation is done at *t = 0*, then

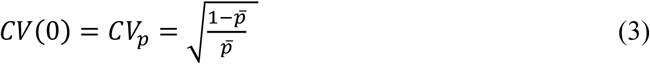

is consistent with the Bernoulli reactivation outcome for the initial cell. Considering 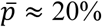 responsive cells in the original population, this would correspond to *CV*(0) ≈ 2. As expected, colony-to-colony fluctuations are high when the reactivation is done early in the lineage expansion as the memory of the initial cell is retained. Over time, these fluctuations decay to zero with each colony equilibrating to the same steady-state % reactivation 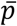. For a fixed time of TNF-α exposure, colony-to-colony fluctuations are enhanced by slower switching between states (Fig. 3B and 3C).

For fitting the predicted *CV* to data, we modify (2)

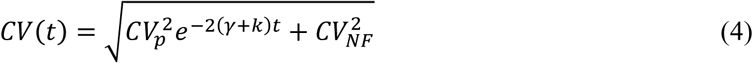

to include a noise floor 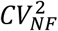 as quantified using fluctuations in % reactivation across unsorted populations. Given an a-priori knowledge of 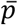, fitting (4) to experimentally measured colony-to-colony fluctuations at different times of exposure infers the decay rate *k* + γ. Note from (1) that 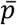 itself is determined by the ratio of *k* and γ. Thus, combining knowledge of both 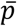 with the inferred decay rate allows both switching rates to be uniquely estimated. Before fitting (4) to the measured colony-to-colony fluctuations, recall that the average % reactivation of single-cell colonies was significantly reduced at week 11 and week 13 (Fig. 2D). For this reason the measured CV in % reactivation at these time points is likely higher than if the mean % reactivation had remained constant at 20%. Assuming CV^2^ to be inversely proportional to the mean % reactivation (as in a binomial distribution), we readjust the CV values at weeks 11 and 13 assuming a 20% mean reactivation. An alternative approach is to simply exclude data points at week 11 and 13, and both approaches yield statistically similar parameter estimates. Fitting model-predicted CV to measured fluctuations reveals the noise decay rate *k* + γ to be 0.24 ± 0.1 per week (Fig. 3D) where the ± denotes the 95% confidence interval as estimated using bootstrapping. Based on a 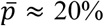, this corresponds to single cells being in the response state for 6.3 ± 2.6 weeks. A key assumption in this calculation is that sorted single-cells can instantly start proliferating. However, the stress of single-cell sorting can create a lag phase before sorted cells can execute normal cellular proliferation. A quick analysis based on cell densities at different times of the colony expansion points to a lag phase of around 2-3 weeks (see details in the Supplementary Information), and accounting for this delay reduces the average time cells stay in the response state to be 2.9 ± 1.2 weeks. Given that only 20% of cells are TNF-α responsive, yields the average time a cell is in the irresponsive state to be 12 ± 4.5 weeks.

## Discussion

The Luria-Delbrück experiment, also called the “Fluctuation Test”, introduced over 75 years ago, demonstrated that genetic mutations arise randomly in the absence of selection – rather than in response to selection – and led to a Nobel Prize (29). Combining this classical fluctuation test with mathematical modeling we have developed a novel methodology to infer timescales of reversible switching between cellular states that exist within the same clonal population. In essence, slower switching rates lead to higher colony-to-colony fluctuations in the assayed phenotype (for example, the % HIV-1 reactivation across colonies) (Figs. 3B and 3C). The key advantage of this method is that it is general enough to be applied to any proliferating cell type, and only involves making a single endpoint measurement. This is especially important for scenarios where a measurement involves killing the cell, and hence the state of the same cell cannot be measured at different time points. Along these lines, the fluctuation test has had tremendous recent success in deciphering stochastic transitions between drug-tolerant states underlying cancer drug resistance, where drug exposure leads to heterogeneous single-cell responses between cell death and survival (30, 31).

We leveraged the fluctuation test to address a fundamental question in HIV-1 latency: is the reactivation of the latent provirus upon LRA exposure purely stochastic or a deterministic function of an underlying cell state (Fig. 1D). Using the Jurkat T-cell line model of HIV-1 latency (JLat 9.2), containing a modified full-length provirus that expresses GFP upon reactivation (Fig. 1B–1C) (25), the fluctuation data strongly points to the heritable model – cells switch between responsive and irresponsive states, and the cell state at the time of TNF-α exposure determines the all-or-none reactivation decision. Remarkably, these states are transiently inherited across generations with a time scale of months (Fig. 3D). This point is exemplified by the fact that even after 5 weeks of clonal expansion there are significant colony-to-colony differences in TNF-α responsiveness (Fig, 1F), and these differences attenuate over time to baseline levels at ≈10 weeks (Fig. 2D). Despite focusing on JLat 9.2, a single JLat clonal integration site, we predict that the heritable model is conserved across integrations but with different time scales (32). The long timescales of switching involved here imply epigenetic mechanisms at play, and indeed, prior work has shown that DNA methylation patterns at the HIV promoter critically regulate provirus reactivation (33-35). Moreover, RNA-induced epigenetic silencing using promoter-targeted shRNAs have been shown to suppress HIV-1 reactivation from latency in JLat 9.2 cells (36). Interestingly, recently constructed high-resolution maps of histone marks in human cells reveal stochastic switching between fully methylated and unmethylated states of DNA at several gene loci (37). Taken together, these results suggest that dynamic, but slow turnovers of epigenetic signatures at the HIV-1 promoter may provide a mechanism for switching between LRA-responsive and irresponsive states.

The measured colony-to-colony fluctuations in the fraction of reactivating cells over time reveals that cells can remain irresponsive to TNF-α for several months before switching to a responsive state. Cells in the responsive state return to become irresponsive after a few weeks. This result has important implications for designing therapies to purge the latent reservoir (9, 38, 39) and suggests that a strategy with multiple LRA treatments spaced over several months may be more effective than single-round, single-LRA treatments (17, 40, 41). Consistent with the idea, previous work has shown that periodic treatments can reactivate more latent virus (40). While this contribution has focused on HIV-1 reactivation in Jurkat T-cells using TNF-α induction, it raises intriguing questions on the generality of transient memory states in the context of other LRAs and cell types. For example, fluctuation-based assays can be done at different proviral integration sites in Jurkat T-Cells, at different TNF-α dosage, in other cell types (for example, myeloid models of HIV-1 latency), and most importantly, using a variety of other LRAs, such as HDAC inhibitors (8, 39, 42), synergistic combinations of PKC agonists (43), DNA cytosine methylation inhibitor 5-aza-2′deoxycytidine (33, 34), and many others reported in the literature. Such studies will be essential to decipher cell-state signatures in the latent reservoir that will guide efforts for HIV-1 treatment. Alternatively, the finding that cells can remain irresponsive to TNF-α for several months before switching to a responsive state may also have implications for additional treatment strategies. The “block and lock” strategy involves silencing HIV-1 expression into a prolonged latent state (16, 44). If the responsiveness of HIV-1 to silencing drugs is governed by the same principle, the responsiveness time scale quantified here may guide the treatment design needed for latency promoting agent administration, in both composition and timing.

## Acknowledgement

AS acknowledges support from NIH grants 5R01GM124446 and 5R01GM126557. YL and RDD acknowledge support from NIH K22 AI120746 and NSF CAREER 1943740.

## Materials and Methods

### Bulk Cell Culture

JLat 9.2 cell line was obtained from the NIH AIDS Reagent Program. Bulk JLat 9.2 cells were cultured in Corning RPMI 1640 w/ L-glutamine and phenol red, with 10% fetal bovine serum (FBS) and 1% penicillin-streptomycin added. Cells were kept in T25 flasks and were passaged twice each week, with a dilution ratio of cell culture to fresh media of about 1:5.

### FACS and JLat Cell Outgrowth Assay

One day before sorting, the concentration of the bulk JLat 9.2 culture was calculated using hemocytometer, and diluted to about 8 × 10^5^ cells/ml. On the day of sorting, four V-bottom 96-well plates were prepared with 100 μl of culture media (same as the media used in bulk cell culture) and 100 μl of pure FBS. During the sorting, one cell was deposited into each well if it passed through the live gate. The plates were subsequently centrifuged to allow cells settle down to the bottom of the wells, and incubated at 37 °C, 5% CO_2_. Plates were checked twice a week, and the contents of wells that turned yellow (indicates cell growth) were transferred to a 48-well plate with 1 ml of media added. After sufficient growth in 48-well plates, cells were expanded into 24-well plates with 2 ml of media, and then 6-well plates after another round of growth. See figure S1 for gating strategy during cell sorting.

### Culturing Colonies Grown From Single-cells

After sufficient growth from sorted single cells, each colony was kept in culture in a 6-well plate, with 3 ml of total culture volume. Cells were passaged twice each week, with a dilution ratio of cell culture to fresh media of about 1:5.

### Latency Reactivation Assay

JLat reactivation assays were conducted once every two weeks, from week 5 to week 13 post-sorting. For each colony, 1 ml of cell suspension was transferred to a well on a flat bottom 24-well plate. TNF-α was added to each well to achieve a final concentration of 10 ng/ml (27, 28). Afterwards cells were incubated at 37 °C, 5% CO_2_ for 24 hours. Samples ran through a BD LSR Fortessa Flow Cytometry Analyzer, and data obtained was subsequently analyzed with FCS Express software. See figure S2 for gating strategy.

### Statistics and Testing

We resorted to bootstrapping by creating re-samplings of the same size as the number of samples between the two sets of data under comparison. We conducted 1,000 unique re-samplings, calculating a CV and mean value for each. We counted the number of times when the re-sampled CV value of a given week is smaller than the control, and when the re-sampled mean value of a given week is larger than the control, and divide the counts by 1,000 to obtain the CV and mean p-values.

A similar procedure is used to generate standard errors that are represented as error bars in Fig. 2D. For each week’s reactivation percentage data, 1,000 unique re-samplings were carried out. The standard error of a given statistic *θ* is calculated as the following.

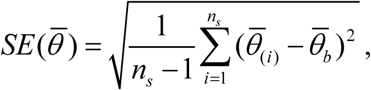

where *n*_*s*_ is the number of unique re-samplings, 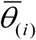 is the mean of the *i*^th^ re-sample, and 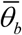 is the mean of all 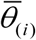.

### Mathematical Modeling

If a cell is in a responsive state at time *t*, i.e., *p*(*t*) = 1 then the probability of it switching to an irresponsive state in the infinitesimal time interval (*t, t* + *dt*) is given by

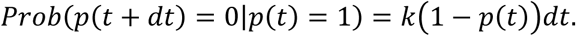

Similarly, the probability of a switch from an irresponsive to responsive state in the infinitesimal time interval (*t, t* + *dt*) is given by

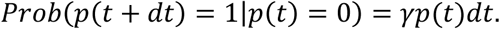

To obtain the time evolution of the first- and second-order statistical moments of *p*(*t*) we use standard tools from moment dynamics that yield (45, 46)

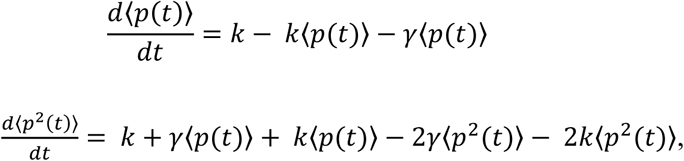

and solving these differential equations at equilibrium result in the steady-state moments given by (1). For the fluctuation test, we assume that a single cell is chosen from the original population with

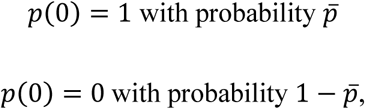

where 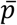 is % reactivation of the original unsorted population. Given this initial random cell state, the fraction of responders *f*(*t*) in the population after time *t* is obtained by solving the mean dynamics of *p*(*t*) leading to

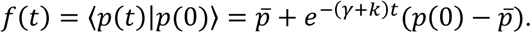

As expected, the average % reactivation across colonies

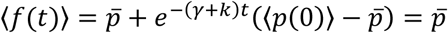

is the same as the average % reactivation of the original unsorted population. The coefficient of variation *CV* in the % reactivation is given by

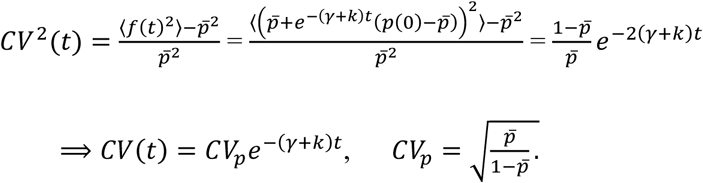

## Notes

### Competing Interest Statement

The authors have declared no competing interest.

