## Supplementary Information for "“A transient heritable memory regulates HIV reactivation from latency”"

### Estimation of growth delay induced by single-cell sorting

It has been observed that using FACS to separate single cells induces stress and delays cell growth, which may contribute to the long memory timescale observed in our experiment. Here we provide an estimation of the cell growth delay from literature data. The doubling time of Jurkat cell line has been measured as around 20.7 hours (1), and has been reported to reach a saturation density of $6 \times 10^6$ cells/ml (2). Cells used in our experiments were kept in exponential growth, combined with the assumption that exponential growth stops at half of the saturation density, it would take a healthy Jurkat cell culture about 23.1 doublings, corresponding to 478.2 hours or 2.85 weeks of growth, to expand from 1 cell to $9 \times 10^6$ cells, the maximum amount of cell in 3 ml of culture under exponential growth.

For the single-cell colonies used in our experiment, the first 14 colonies reached saturation density at about 4.5 weeks, while the rest 2 colonies reached saturation density no later than 6.5 weeks. Assuming the single cells exhibit all-or-nothing growth behaviors, stopping their growth at first before resuming normal growth with doubling time of 20.7 hours, this would correspond to a growth delay of 1.65 – 3.65 weeks.

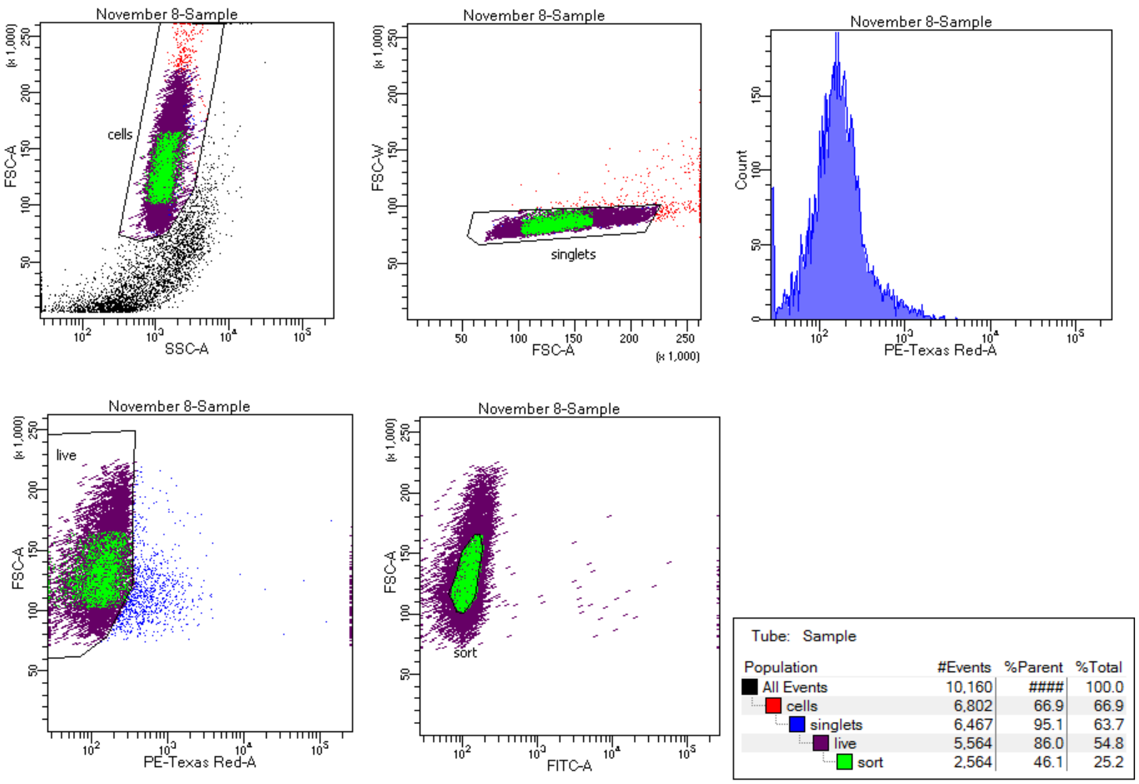


**Figure S1. Gating strategy during single-cell sorting of JLat 9.2.** Propidium iodide (PI) staining was used to distinguish live cells from dead ones. Four layers of gating were implemented during the sorting. The first layer ("cells" gate) gets rid of debris and dead cells. The second layer ("singlets" gate) removes large particles and ensures every cell passes through is a singlet. The third layer ("live" gate) selects cells with low fluorescence in the PE-Texas Red channel, which captures PI fluorescence, and removes dead cells. The final gate ("sort" gate) only includes the center of the live cell distribution and selects cells that are representative of the population behavior. The cells in the "sort" gate were plated onto 96-well plates, with one cell in each well.

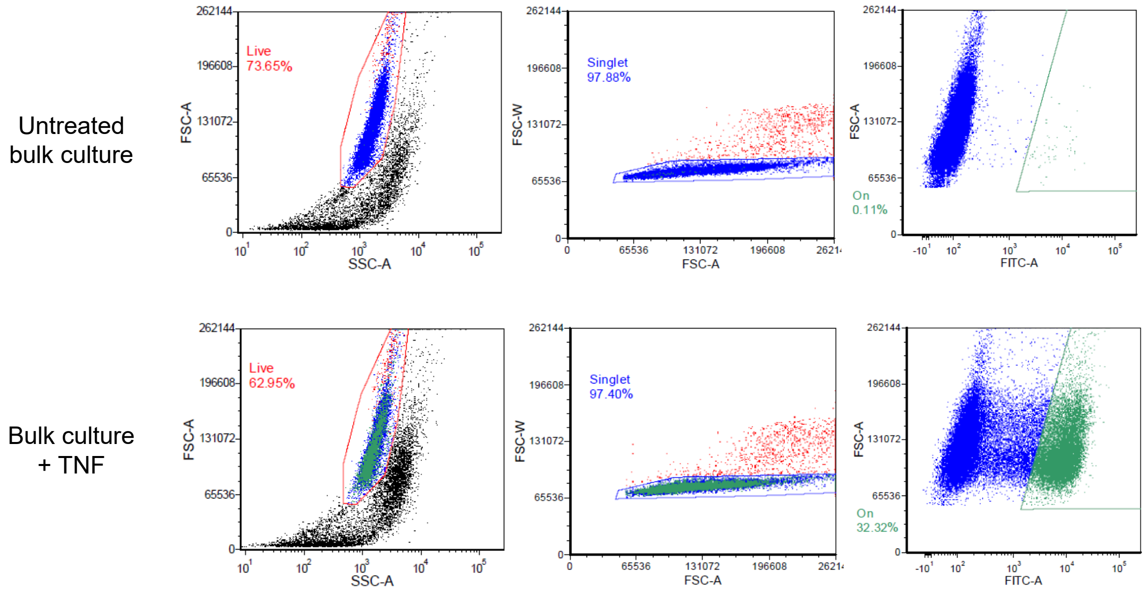


**Figure S2. Gating strategy for latency reversal assay.** We used a three-layered gating to quantify the reactivation percentage of JLat 9.2. Shown here is a sample gating for untreated bulk culture (top row) and bulk culture after 24 hours of TNF treatment (bottom row). The first layer ("Live" gate) gets rid of debris and dead cells. The second layer ("Singlet" gate) gets rid of large cells that are potentially doublets. The third layer ("On" gate) only includes cells with significantly higher fluorescence in FITC channel, which captures GFP fluorescence. The ratio of number of cells in the "On" gate (green dots) over the number of all cells in the "Singlet" gate (green + blue dots) were used as the quantification of reactivation percentage.
